# Bleaching-independent, whole-cell, 3D and multi-color STED imaging with exchangeable fluorophores

**DOI:** 10.1101/420638

**Authors:** Christoph Spahn, Jonathan B. Grimm, Luke D. Lavis, Marko Lampe, Mike Heilemann

## Abstract

We demonstrate bleaching-independent STED microscopy using fluorogenic labels that reversibly bind to their target structure. A constant exchange of labels guarantees the removal of photobleached fluorophores and their replacement by intact fluorophores, thereby circumventing bleaching-related limitations of STED super-resolution imaging in fixed and living cells. Foremost, we achieve a constant labeling density and demonstrate a fluorescence signal for long and theoretically unlimited acquisition times. Using this concept, we demonstrate whole-cell, 3D, multi-color and live cell STED microscopy with up to 100 min acquisition time.

## Main

Fluorophore bleaching is a fundamental challenge in super-resolution STED microscopy. Minimizing photobleaching comes with more fluorophores that contribute to signal-to-noise and thus resolution. This is particularly beneficial for super-resolution imaging, especially in live-cell and 3D imaging. In the context of STED microscopy ^1,2^, this challenge was addressed, for example, by developing optical schemes that dynamically tune the excitation light during image acquisition ^3^, or by developing photostable fluorophores (reviewed in Zheng & Lavis ^4^). Finding solutions to bypass fluorophore bleaching can advance STED microscopy towards whole-cell imaging, since bleaching of out-of-focus fluorophores is reduced and a large-volume imaging can be achieved. Another important application of reduced photobleaching is long-term live cell STED imaging, as shown recently for STED imaging of extracellular space of living brain slices ^5^.

In single-molecule localization microscopy ^6^, photobleaching can be bypassed using fluorophores that reversibly bind to their target structure, a concept that originally was realized in Point Accumulation for Imaging in Nanoscale Topography (PAINT) ^7^. We repurposed this idea, and introduce bleaching-independent STED (biSTED) microscopy by using fluorophores that reversibly bind to and dynamically exchange from their target structure (Figure 1a). This procedure efficiently replaces fluorophores that were imaged and eventually photobleached by new and intact fluorophores. This concept requires fast exchange kinetics of the fluorophores and a sufficiently large reservoir of unbleached (intact) fluorophores. It must also ensure high-density labeling of the target structure.

**Figure 1:**
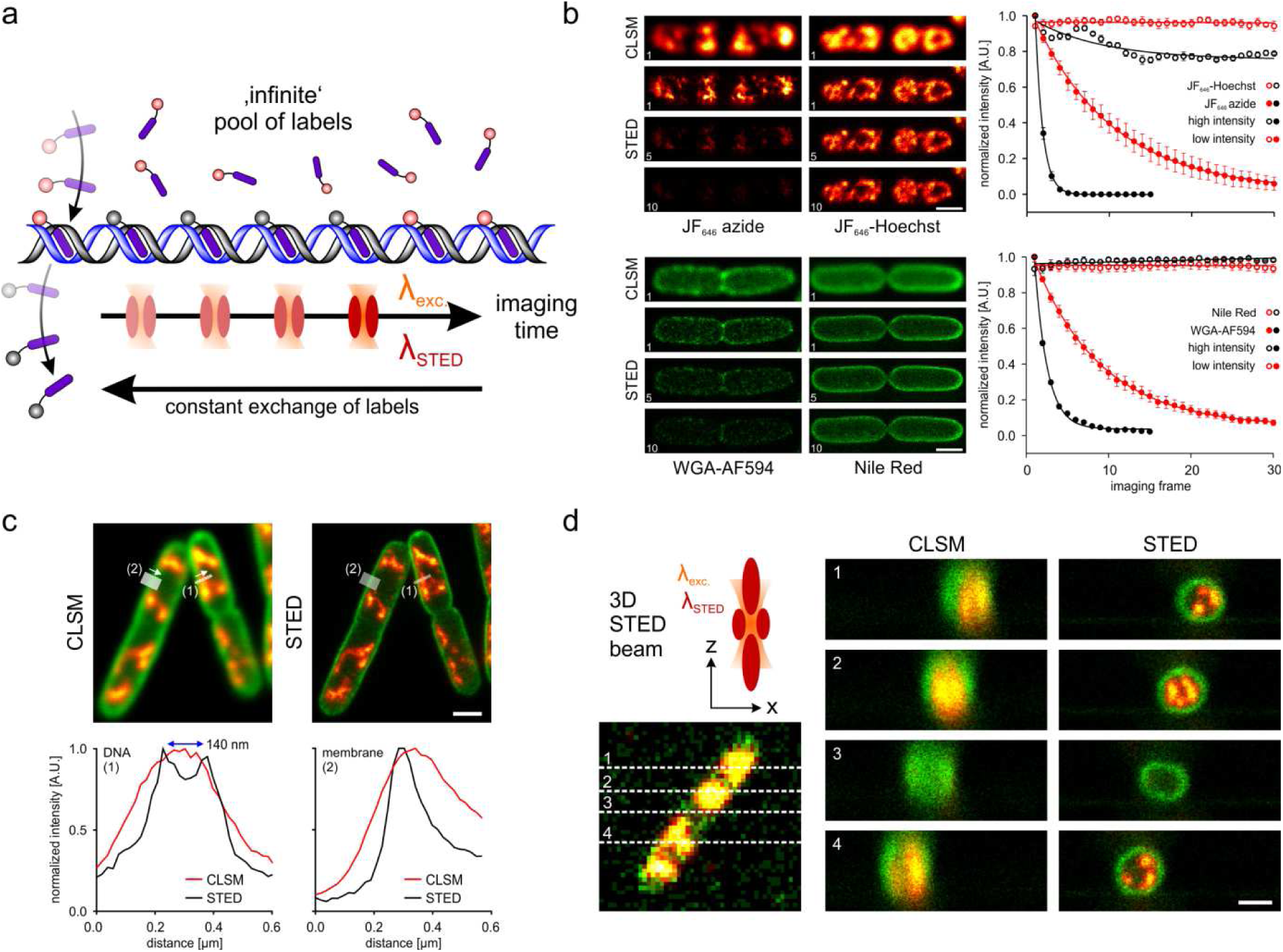
Bleaching-independent STED microscopy. **(a)** Fluorophores that dynamically bind and unbind to their target structure ensure a constant exchange with a quasi-infinite pool, compensating for irreversible photobleaching and allowing for long-term STED imaging. **(b)** Multi-frame STED microscopy comparing stationary and reversibly binding labels targeting the *E. coli* membrane or nucleoid. Cells were labeled using either a static (left panel) or dynamic labeling approach (right panel). Static labeling was achieved using click-chemistry (DNA, red hot) or sugar-binding lectins (cell wall, green). A high concentration of the exchangeable labels JF_646_-Hoechst and Nile Red were applied for dynamic labeling of DNA (red hot) and *E. coli* membranes (green), respectively. Fluorescence intensities were extracted for each imaging frame (data points represent mean values of at least 3 cells and error bars the respective standard deviation) and fitted with a mono-exponential decay. **(c)** Parallel two-color STED of *E. coli* DNA (red hot) and membrane (green) using one depletion wavelength at 775 nm. Both cross-sections of DNA filaments (1) and membrane (2) illustrate the resolution enhancement provided by 2D STED. **(d)** Two-color 3D STED imaging with near isotropic resolution. The cell shown in the low-resolution XY-overview image was measured in XZ-scanning mode. Dashed lines indicate the position of XZ cross-sections shown in the right image panel (the full stack is provided as Supplementary Video 1). Scale bars are 1 μm.

We demonstrate bleaching-independent STED by implementing exchangeable fluorophores that were previously used in PAINT imaging ^7^. We first investigated whether fluorogenic PAINT fluorophores can in principle be used for STED imaging, using the membrane stain Nile Red ^7^ and the DNA stain JF_646_-Hoechst ^8^. We demonstrate sub-diffraction spatial resolution STED imaging with these exchangeable fluorophores in *E. coli* cells (Figure 1b). In contrast to single-molecule PAINT imaging, where a low labeling density ensures the detection of single emitters, our concept requires higher concentrations of fluorophores in the buffer (~100 nM – 1 μM) in order to ensure a constantly high labeling density despite the exchange dynamics.

Next, we investigated whether a constant and dynamic exchange of fluorophores occurs, which should enable multiple rounds of STED imaging of the same cell without any loss of fluorescence signal. For this purpose, we again chose *E. coli* cells as cellular test system for imaging, because of their small size and regular shape that facilitates quantitative intensity analysis. We recorded a sequence of STED images of the bacterial cell envelope using either the exchangeable label Nile Red, or the stationary label Alexa-Fluor 594-conjugated wheat germ agglutinin (WGA-AF594). The quantification of the fluorescence intensity over multiple imaging frames showed photobleaching for the stationary label, WGA-AF594, and a constant fluorescence signal for the exchangeable label, Nile Red (Figure 1b). We next recorded a sequence of STED images of the bacterial nucleoid, using either the exchangeable label JF_646_-Hoechst, or the same fluorophore, JF646, as stationary label targeting DNA through click labeling of EdU-treated cells ^9^. Again, we observed photobleaching for the stationary label and a constant fluorescence signal for the exchangeable label (Figure 1b). Under similar imaging conditions, photobleaching was also observed for ATTO647N (Figure S1), a fluorophore that is sufficiently photostable to enable single-molecule STED imaging ^10^. Notably, the labeling density that we observed for the stationary and the exchangeable labels appeared very similar, which demonstrates efficient high-density labeling. Note that the higher concentration of these exchangeable labels in the imaging buffer does not increase the background signal in STED experiments.

We next demonstrated super-resolution imaging of the bacterial nucleoid and envelope in two-color, 2D-STED (Figure 1c), as well as in 2-color, 3D-STED mode (Figure 1d; Supplementary Video 1). Cells were labeled with both exchangeable fluorophores, Nile Red and JF_646_-Hoechst, simultaneously. The 3D STED images clearly reveal the radial positioning of DNA (red hot) in proximity of the bacterial plasma membrane (green) (Figure 1d), potentially mediated by coupling of transcription, translation and membrane insertion (transertion) of proteins ^11^. We note that Nile Red and JF_646_- Hoechst allowed recording STED images using the same depletion laser (emitting at 775 nm), which simplifies the optical setup and image acquisition. Another advantage of these two exchangeable fluorophores is that no specific cellular treatment is needed (e.g. EdU incorporation for high-density labeling of DNA), further simplifying sample preparation and avoiding undesired stress for cells.

We next applied the exchangeable labels Nile Red and JF_646_-Hoechst to STED imaging of eukaryotic cells (Figure 2). First, we demonstrate two-color STED microscopy with sub-diffraction spatial resolution (Figure 2a). The STED images of the Nile Red channel show structural features of various membrane-surrounded cellular organelles, e.g. lipid droplets, mitochondria and the nuclear lamina, which were elusive in the respective CLSM image (Figure 2a).

**Figure 2:**
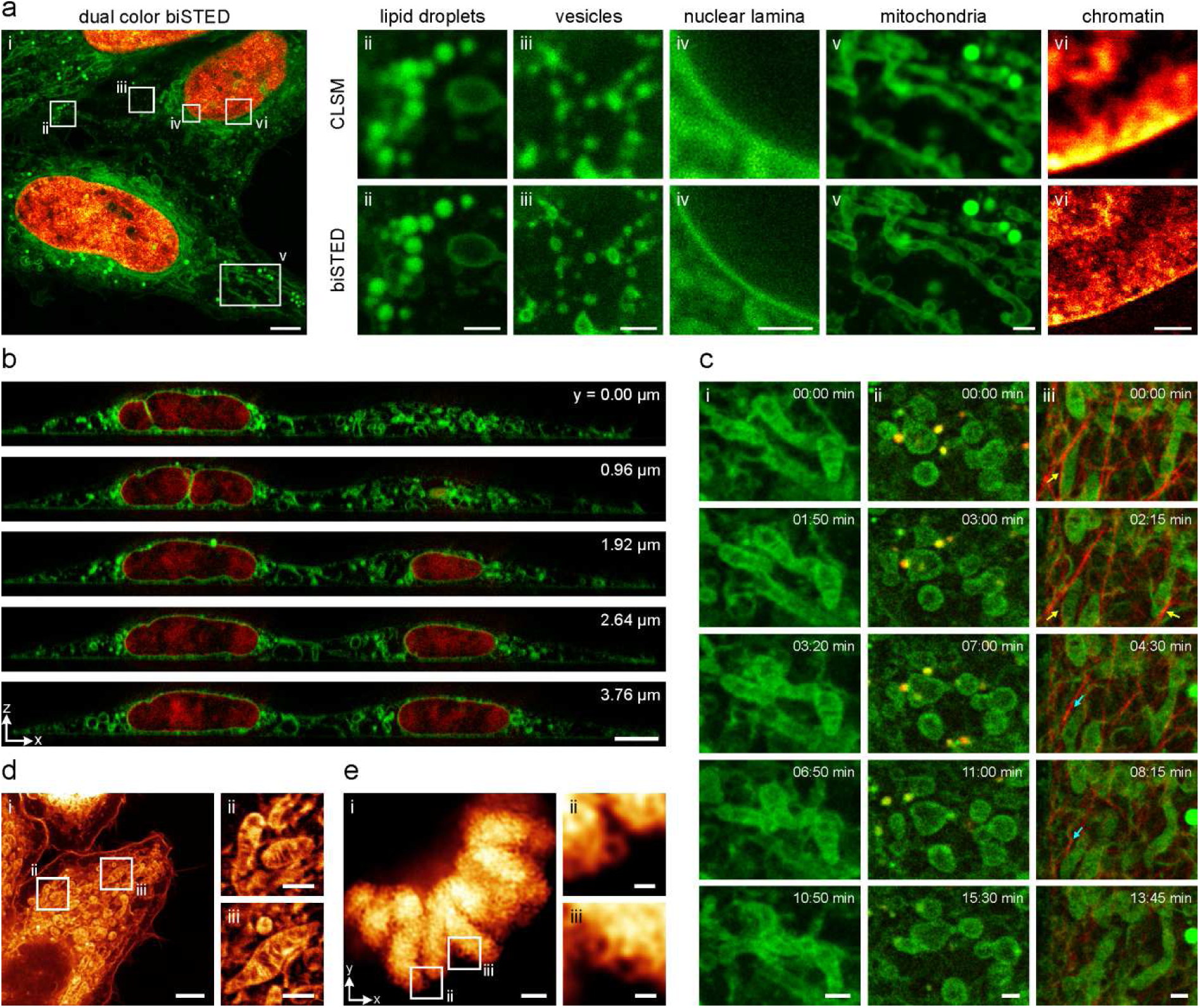
Dual-color 2D and 3D STED imaging in fixed whole cells, and live cell STED imaging using reversibly binding dyes. **(a)** 2-color biSTED image of a fixed HeLa cell (i). DNA (red hot) is labeled with JF_646_-Hoechst and cellular membranes with Nile Red (green). Various membranous structures can be resolved with Nile Red, such as lipid droplets (ii), vesicular structures (iii), the nuclear lamina (iv) and mitochondria (v). Super-resolution of nano-scale DNA features of DNA is achieved with the exchangeable fluorophore JF_646_-Hoechst (vi). **(b)** 3D-STED imaging of fixed HeLa cells labeled for membranes (green) and DNA (red), achieving near isotropic resolution (the comparison to the CLSM image stack is shown in Supplementary Video 2). The image stack was recorded in XZ-scanning mode. **(c)** Live-cell STED imaging with long acquisition times of HeLa cells stained with Nile Red (i), Nile Red and SiR-lysosome (ii) or Nile Red and SiR-tubulin (iii). Arrows in (iii) indicate association (yellow) or dissociation (cyan) of mitochondria and microtubules. **(d)** biSTED image of mitochondria in a fixed HeLa cell, visualized using Nile Red. (i) Average image (raw images) of 4 imaging planes, corresponding to a 240 nm thick section. (ii, iii) Magnified ROIs from (i), showing mitochondrial cristae. **(e)** High-resolution, deconvolved biSTED image of mitotic chromosomes in a fixed HeLa cell, visualized with JF_646_-Hoechst (i). (ii, ii) Magnified regions from (i) showing small chromatin loops. Scale bars are 5 μm (a i, b), 3 μm (e i), 1 μm (a ii-vi, d i, e ii-iii, c) and 250 nm (d ii-iii).

A key advantage of bleaching-independent imaging is that a large number of images at different axial positions can be recorded, without compromising the fluorescence signal in out-of-focus imaging planes by the excitation or depletion laser. To demonstrate this, we recorded whole-cell two-color 3D STED images using a 3D optical depletion scheme in both XZ (Figure 2b, Supplementary Videos 2 and 3) and XY scanning mode (Figure S2), with a constant fluorescence signal over time. Without dynamic fluorophore exchange, the imaging conditions need to consider bleaching as one critical parameter, which demands a compromise in resolution or signal-to-noise ratio. Using exchangeable fluorophores, such optimizations are not necessary anymore.

A big challenge for all super-resolution methods is live cell imaging. Bypassing fluorophore bleaching would allow long-term observation of single cells and following the dynamics of cellular features in an (almost) quantitative manner avoiding misinterpretations due to photobleached labels. Here, we demonstrate time-lapse STED imaging of a living cell labeled with Nile Red (Figure 2c i; Supplementary Video 4), enabling to follow organelle dynamics while displaying no detectable change in average fluorescent intensity (Figure S3). Since Nile Red can be effectively depleted using far-red depletion lasers (here λ_dep_ = 775 nm), it can be combined with other membrane permeable and live cell compatible SiR-conjugates ^12^. This allowed us to perform live-cell dual-color STED imaging of lysosomes (SiR-lysosome), following the dynamics of these with respect to other cellular organelles (Figure 2c ii, Supplementary Video 5). In addition, the tight association of mitochondria and microtubules could be visualized using a tubulin-binding SiR-conjugate (Figure 2c iii, Supplementary Video 6). In contrast to the decreasing signal provided by SiR-tubulin, the Nile Red signal remained constant over the whole STED imaging period (up to 100 min) under imaging conditions providing enhanced spatial resolution (Figure S4, Supplementary Video 7).

Constant label-exchange is not only beneficial for 3D and live-cell measurements. It also allows a higher degree of optimization of imaging parameters for single-color high-resolution STED imaging in fixed specimen. This is limited if static labels are used, due to irreversible photobleaching affecting any STED image recorded with high irradiation intensities. By optimizing imaging parameters for the individual exchangeable labels, we were able to visualize intriguing structural details of mitochondria (Nile Red, Figure 2d, Figure S5) and mitotic chromosomes (JF_646_-Hoechst, Figure 2e). While mitochondrial cristae membranes were previously investigated using electron microscopy (EM) and provided a high level of structural information ^13^, they have to our knowledge not been resolved to a comparable level as shown in this work using fluorescence microscopy. As another example for optimized single-color STED in fixed cells, we visualized the mitotic chromosome in fixed HeLa cells using the exchangeable label JF_646_-Hoechst, followed by deconvolution of the resulting STED image (Figure 2e, Figure S6). The images show a grained chromosome structure (Figure 2e, i) and small DNA loops (Figure 2e, ii and iii). These key features of mitotic chromosomes are also known from EM studies ^14^, but were just recently visualized by fluorescence microscopy using lattice light sheet-PAINT microscopy ^8^.

In conclusion, we present a simple and powerful experimental protocol for bleaching-independent STED microscopy. Our approach builds on exchangeable fluorophores, which are commonly used for single-molecule localization microscopy. We demonstrate that exchangeable labels can achieve high labeling density, circumvent fluorophore bleaching and allow for whole-cell, multi-color and live cell STED imaging. This concept can be extended to other labels that reversibly bind to their target structure, and can be used in correlated light and electron microscopy ^15,16^, which opens many new application windows for STED microscopy.

## Acknowledgment

We thank the Advanced Light Microscopy Facility (ALMF) at the European Molecular Biology Laboratory (EMBL), Leica Microsystems and Abberior Instruments for support. We also thank Petra Freund, George Galea and Juan Jung for help with cell culture and sample preparation. MH acknowledges funding by the German Science Foundation (SFB 902).

## Data availability

Data is available upon request from the authors.

## References

1. Blom, H. & Widengren, J. Stimulated Emission Depletion Microscopy. Chemical reviews 117, 7377–7427 (2017).

2. Vicidomini, G., Bianchini, P. & Diaspro, A. STED super-resolved microscopy. Nature methods 15, 173–182 (2018).

3. Heine, J. et al. Adaptive-illumination STED nanoscopy. Proceedings of the National Academy of Sciences of the United States of America 114, 9797–9802 (2017).

4. Zheng, Q. & Lavis, L.D. Development of photostable fluorophores for molecular imaging. Current opinion in chemical biology 39, 32–38 (2017).

5. Tonnesen, J., Inavalli, V. & Nagerl, U.V. Super-Resolution Imaging of the Extracellular Space in Living Brain Tissue. Cell 172, 1108–1121 e1115 (2018).

6. Sauer, M. & Heilemann, M. Single-Molecule Localization Microscopy in Eukaryotes. Chemical reviews 117, 7478–7509 (2017).

7. Sharonov, A. & Hochstrasser, R.M. Wide-field subdiffraction imaging by accumulated binding of diffusing probes. Proceedings of the National Academy of Sciences of the United States of America 103, 18911–18916 (2006).

8. Legant, W.R. et al. High-density three-dimensional localization microscopy across large volumes. Nature methods 13, 359–365 (2016).

9. Spahn, C., Endesfelder, U. & Heilemann, M. Super-resolution imaging of Escherichia coli nucleoids reveals highly structured and asymmetric segregation during fast growth. Journal of structural biology 185, 243–249 (2014).

10. Kasper, R. et al. Single-molecule STED microscopy with photostable organic fluorophores. Small 6, 1379–1384 (2010).

11. Bakshi, S., Choi, H. & Weisshaar, J.C. The spatial biology of transcription and translation in rapidly growing Escherichia coli. Frontiers in microbiology 6, 636 (2015).

12. Lukinavicius, G. et al. Fluorogenic probes for live-cell imaging of the cytoskeleton. Nature methods 11, 731–733 (2014).

13. Zick, M., Rabl, R. & Reichert, A.S. Cristae formation-linking ultrastructure and function of mitochondria. Biochimica et biophysica acta 1793, 5–19 (2009).

14. Marsden, M.P. & Laemmli, U.K. Metaphase chromosome structure: evidence for a radial loop model. Cell 17, 849–858 (1979).

15. Perkovic, M. et al. Correlative light-and electron microscopy with chemical tags. Journal of structural biology 186, 205–213 (2014).

16. de Souza, N. Super-resolution CLEM. Nature methods 12, 37 (2014).

